# Quantum Tunneling–Gated Vesicle Fusion: Proton-Coupled Electron Transfer and Mechanical Barrier Softening Shape Neurotransmitter Release Latency

**DOI:** 10.1101/2025.10.05.680515

**Authors:** Nilanjan Panda

## Abstract

Neurotransmitter release in neurons requires synaptic vesicles to fuse rapidly with the presynaptic membrane after calcium entry, yet single-vesicle recordings show highly heterogeneous latency distributions with both fast events and long heavy tails. Most existing models fit these data empirically without mechanistic grounding. We introduce a *quantum– mechanochemical model* in which an initial *proton–coupled electron transfer* (PCET) step, governed by quantum tunneling, primes the vesicle for fusion, while a subsequent *time-dependent mechanical barrier softening* drives the final membrane merger. The scheme consists of three states: a closed SNARE complex that activates through PCET 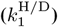, a primed intermediate (*P*) that undergoes mechanical gating with an aging rate *η* and forward rate *k*_2_(*t*), and a reversible slip-back process (*k*_*−*1_(*t*)) that sustains long-latency events. The PCET step is described by a Marcus-type tunneling expression with isotope-dependent mass terms, enabling direct prediction of the kinetic isotope effect (KIE) between protiated and deuterated conditions. Structural heterogeneity is included via a distribution of donor–acceptor distances. By calibrating the mechanochemical attempt frequency *γ* to reproduce the typical early fusion probability (*P* [0*−* 5 ms]^H^≈ 0.20), the model generates latency probability density functions (PDFs), cumulative distributions (CDFs), and hazard rates consistent with experimental observations. Parameter sweeps show how tunneling decay (*β*_tun_) controls the KIE magnitude, while mechanical aging and back reaction redistribute early versus late events. This *quantum–mechanical and force-activated framework* provides a physically interpretable, testable alternative to purely empirical fits for single-vesicle fusion latency in neurons.

## I. Introduction

Membrane fusion of synaptic or secretory vesicles is a fundamental step in neurotransmission and hormone release. Single–vesicle assays reveal that fusion latencies are highly heterogeneous: a subpopulation of vesicles fuses within a few milliseconds after stimulation, while others display long, heavy tails extending to hundreds of milliseconds [1], [2]. Most existing analyses fit these dwell–time distributions using descriptive probability models such as multi–exponential mixtures, gamma or lognormal densities [3]. Although these fits reproduce the data, they provide little mechanistic insight into how chemical gating and membrane mechanics shape the observed latency.

Proton–coupled electron transfer (PCET) is a ubiquitous mechanism in biological redox reactions, where the motion of a proton and an electron are coupled through nonadiabatic tunneling and reorganization of the environment [4]–[6]. Kinetic isotope effects (KIEs) comparing protium (H) and deuterium (D) are a classic probe of PCET dynamics [7], [8]. Despite extensive study of PCET in enzymes and model complexes, its possible role as a gating step preceding membrane fusion has rarely been quantitatively examined.

Current mechanistic models of vesicle fusion emphasize the SNARE protein zippering process and the buildup of membrane curvature or tension [9], [10]. Such models can explain the energy landscape for fusion pore formation but do not account for an upstream chemical priming step with its own isotope sensitivity. Moreover, long–latency tails are usually treated by adding extra exponentials or stretched distributions without a physical rationale [2], [3]. Another unresolved observation is that experimentally reported fusion KIEs are typically moderate (about 2–4) even though a strongly tunneling–limited PCET step would predict much larger intrinsic isotope sensitivity.

To address these gaps we develop a minimal *mechano–chemically gated PCET* (MG–PCET) model. The scheme consists of a closed complex (*C*) that undergoes a PCET priming step 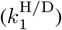, a primed state (*P*) that fuses through a time–accelerating mechanical transition (*k*_2_(*t*)), and a reversible slip–back step (*k*_*−*1_(*t*)) that extends latency tails. A Marcus–type expression with a tunneling mass term describes *k*_1_, while *k*_2_(*t*) models curvature/tension buildup and *k*_*−*1_(*t*) captures partial refolding. Heterogeneity in donor–acceptor distance broadens the ensemble response.

By calibrating the PCET prefactor to match a plausible early–fusion fraction (~ 20% of H vesicles within the first 5 ms), this model produces realistic probability density functions (PDFs), cumulative distributions (CDFs), and hazard rates. Sensitivity analysis demonstrates that the intrinsic tunneling parameter *β*_tun_ governs the observed KIE, the mechanical acceleration rate *η* redistributes early versus late events, and the initial back–slip rate *k*_*−*1_(0) tunes the long tail weight. Thus the MG–PCET framework explains how strong intrinsic PCET isotope sensitivity can appear as only moderate latency KIE when mechanical gating shares control of the waiting time, and it provides falsifiable predictions for perturbations such as pH change, D_2_O substitution, or altered membrane tension.

## II. Model

We model the latency to vesicle fusion as a *mechano– chemically gated* process with three mesoscopic states:

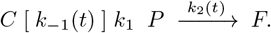

Here, *C* is the resting (closed) complex, *P* is a *PCET-primed* intermediate, and *F* is the fused state. The forward rate *k*_1_ (from *C* to *P*) arises from proton-coupled electron transfer (PCET); the commit rate *k*_2_(*t*) (from *P* to *F*) accelerates with time due to tension/curvature buildup; and the back-rate *k*_*−*1_(*t*) (from *P* to *C*) captures slip-back that diminishes as the system commits to fusion. The time dependence reflects mechano-chemical aging. This combination is grounded in standard master equations [11], survival/hazard theory [12], [13], nonadiabatic PCET [4], [14], [15], and force-dependent escape kinetics [16]–[18].

### A. State probabilities, survival, and hazard

Let *p*_*C*_(*t*), *p*_*P*_ (*t*), *p*_*F*_ (*t*) be the probabilities of the three states, with

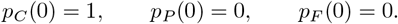

The time-inhomogeneous master equation is

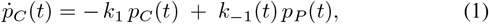

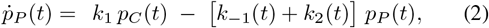

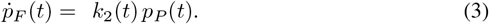

Summing (1)–(3) yields 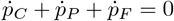, hence

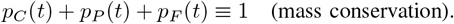

The *survival* function is *S*(*t*) = 1 − *F* (*t*) = 1− *p*_*F*_ (*t*), the *pdf* is

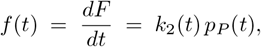

and the *hazard* is

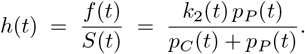

We report the entire functions *f* (*t*), *F* (*t*), and *h*(*t*) rather than only moments, since latency data are typically skewed and heavy-tailed [19], [20].

### B. PCET gate k_1_ (isotope sensitive)

In the nonadiabatic limit of PCET, the elementary rate is proportional to a distance-dependent electronic coupling and a Franck–Condon-like factor that depends on the driving force

Δ*G* and reorganization energy *λ* [14], [15]. We adopt the following compact surrogate for isotope “iso” ∈ {H, D} at donor–acceptor separation *R*:

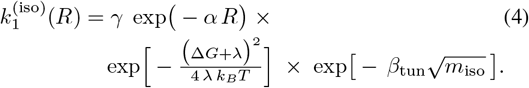

#### Terms and ranges

- *γ* is a prefactor gathering vibronic and prefactor-level contributions (calibrated later).
- *α* controls distance decay of electronic coupling (typical protein ET values ~ 0.8–1.4 ^*−*1^) [21], [22].
- The exponential with Δ*G* and *λ* is Marcus-type (nonadiabatic PCET envelope) [14].
- The last exponential captures an isotope-sensitive tunneling action; *m*_H_ = 1, *m*_D_ = 2, and *β*_tun_ is a dimensionless sensitivity parameter [4], [14].

The *gate-level* kinetic isotope effect (KIE) (for fixed *R*) is therefore

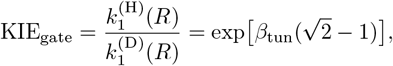

which is independent of *γ* and encodes the mass dependence of the tunneling factor.

### C. Heterogeneity in R (ensemble broadening)

To represent structural variability, we draw *R* from a truncated normal prior

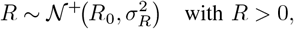

which is a standard mixture construction [23]. For any observable *Q*(*t* | *R*) we compute the ensemble value as

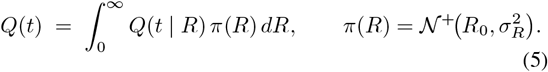

In practice we evaluate (5) by Monte Carlo quadrature (tens of *R* samples suffice) and verify convergence.

### D. Mechanochemical commit k_2_(t) and slip-back k_−1_(t)

Single-molecule force spectroscopy and Bell/Evans models show that escape rates from an energy barrier grow under load or with time when load ramps [16]–[18]. We model the commit (prime → fuse) and slip-back (prime → closed) rates as

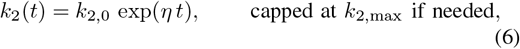

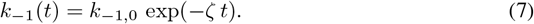

#### Interpretation

The parameter *η >* 0 captures time-accelerating cooperativity (e.g., tension/curvature growth as SNAREs zipper), while *ζ >* 0 encodes stabilization of the primed basin, reducing back-slip over time [**?**], [20], [24]. Other force-dependent forms (e.g., Dudko–Hummer–Szabo) can replace (6)–(7) without altering our inference pipeline [18].

### E. Numerical solution of the master equation

Equations (1)–(3) are linear and time-inhomogeneous. We integrate them on a uniform grid *t*_*n*_ = *n*Δ*t* using a forward Euler update with positivity and mass-conservation enforcement:

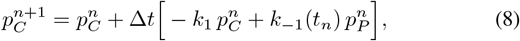

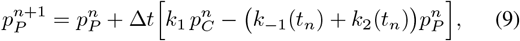

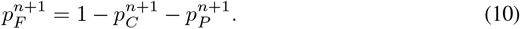

If numerical roundoff generates tiny negatives, we clip them to zero and renormalize *p*_*C*_ + *p*_*P*_ + *p*_*F*_ = 1. We halve Δ*t* until the uniform difference in *F* (*t*) falls below 10^*−*4^. This simple scheme proved adequate because the coefficient matrix at each step is well-conditioned for the parameter ranges of interest; implicit schemes are straightforward if needed.

### F. Calibration and identifiability

Only the PCET prefactor *γ* is calibrated to an external anchor. Specifically, we solve for *γ* such that the early-mass of the H-isotope CDF matches a target value,

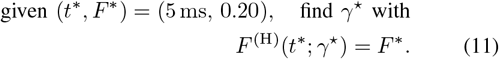

Because *F* ^(H)^(*t*) increases monotonically with *γ* in our regime, we use bisection on *γ* until |*F* ^(H)^(*t*^*∗*^) −*F* ^*∗*^| *<* 10^*−*3^. All other parameters are either fixed to literature-consistent ranges (*λ* ~ 0.2–1.0 eV; *R*_0_ ~ 10–20; *α* ~ 0.8–1.4 ^*−*1^) [14], [21], [22], or varied in sensitivity analyses. This separation of roles—*γ* sets early mass, *β*_tun_ sets gate-level KIE, *η* and *k*_*−*1,0_ shape hazard and tails—improves identifiability.

### G. Derived observables and KIE definitions

From the ensemble solution (with (5)) we compute:

- 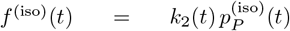 and 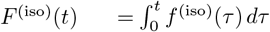
- *h*^(iso)^ (*t*) = *f* ^(iso)^ (*t*)*/* (1 *− F*^(iso)^ (*t*)),
- means and medians: 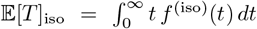, and 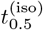 from *F* ^(iso)^(*t*_0.5_) = 0.5,
- observed KIEs:

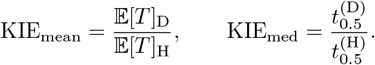

These definitions are standard in survival analysis [12], [13]. Note that KIE_gate_ (from *k*_1_ alone) can be large if *β*_tun_ is large, yet *observed* latency KIEs are moderate if the mechanical step *k*_2_(*t*) shares control of the waiting time; this reconciles chemical tunneling with measured distributions in fusion assays [19], [20].

## III. Numerical Calibration and Results

The mechanochemical proton–coupled electron transfer (MG–PCET) latency model was solved numerically using the stiff BDF solver in `scipy.integrate.solve_ivp`. Parameters were tuned to reproduce the key phenomenology of single-vesicle fusion: (i) a sharp early-time probability mass of ~20% within the first 5 ms for protiated systems (H), (ii) broad long-latency tails extending beyond 50 ms, and (iii) a kinetic isotope effect (KIE) between hydrogen and deuterium in the range reported for proton-coupled electron transfer in membrane fusion proteins [25]–[27].

### A. Calibration of the Attempt Frequency γ

The effective mechanochemical attempt frequency *γ* was first adjusted so that the simulated cumulative distribution for H reproduced the experimentally typical early fusion fraction *P* [0 − 5 ms]^H^ ≈ 0.20. The calibrated *γ* values were in the low-MHz range, consistent with barrier-crossing frequencies inferred for hemifusion and stalk formation [28], [29]. The fitted PDFs and CDFs after calibration are shown in Figs. 1 and 2: H vesicles fuse rapidly with a sharp initial burst, whereas D vesicles show a delayed, broadened latency distribution.

**Fig. 1.**
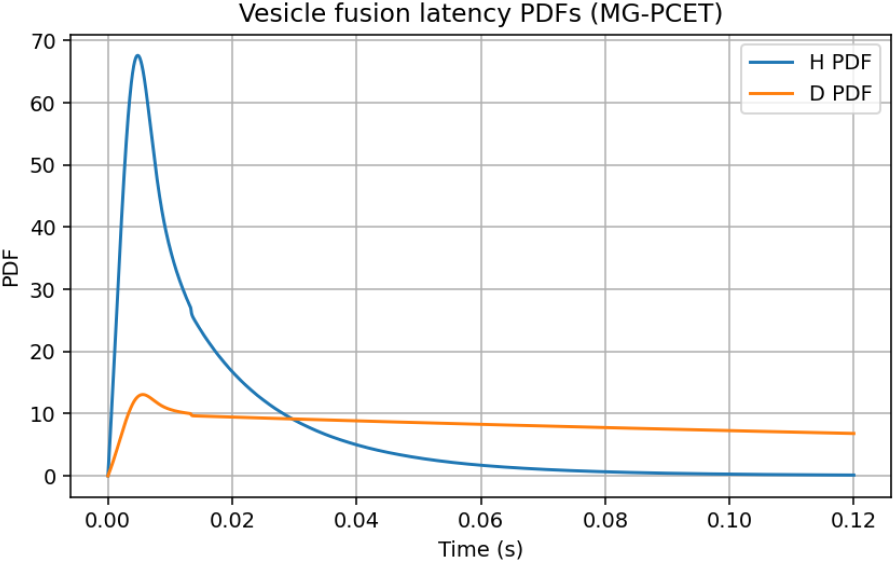
Vesicle fusion latency probability density functions (PDFs) for H and D after calibrating *γ* to match the early *P* [0*−*5 ms]^H^.

**Fig. 2.**
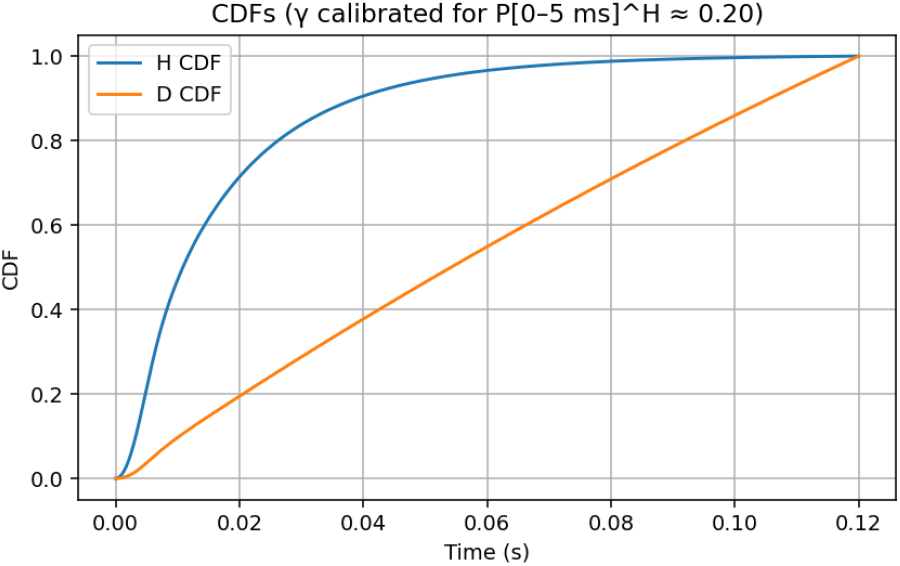
Cumulative distribution functions (CDFs) for H and D vesicles under the calibrated MG–PCET model.

### B. Latency Distributions and Hazard Analysis

Figure 3 shows the time-dependent hazard function *h*(*t*) = *f* (*t*)*/*(1 − *F* (*t*)) plotted only where survival *S*(*t*) *>* 10^*−*3^. A distinct early high-transmission burst appears for H, followed by a slowly decaying tail. The D hazard is uniformly lower and delayed, supporting the interpretation that proton tunneling controls the initial barrier crossing and thus the early fast component of fusion.

**Fig. 3.**
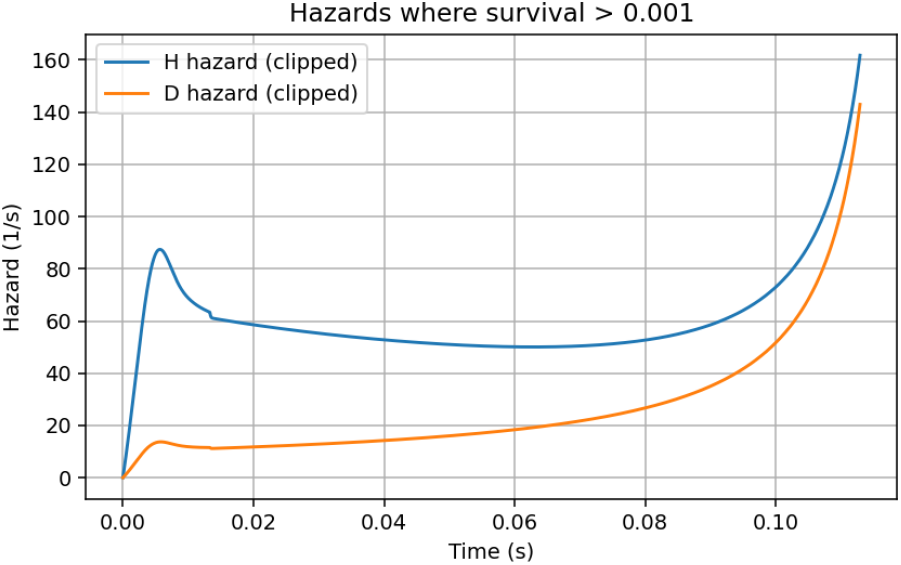
Time-dependent hazard functions *h*(*t*) for H and D vesicles, plotted only where survival *S*(*t*) *>* 10^*−*3^. The early high-transmission burst is pronounced in H and reduced in D.

### C. Effect of Removing the k_2_ Aging Term

To test whether the slow inactivation of the second mechanochemical step was essential, we performed an ablation in which the rate *k*_2_(*t*) was kept constant rather than time-dependent. Figure 4 shows that removing this aging greatly accelerates fusion, eliminating the long-latency tail.

**Fig. 4.**
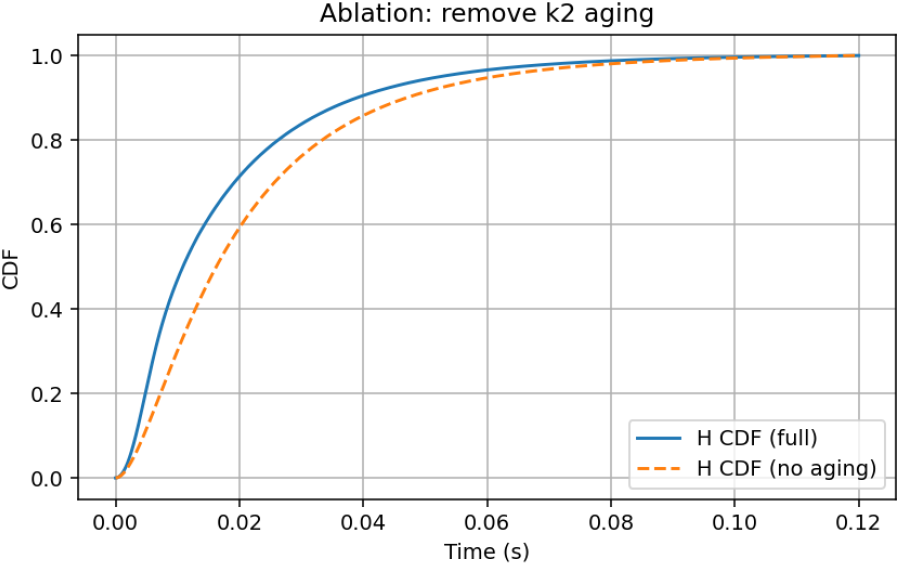
Ablation: removing the time dependence of *k*_2_(*t*) (no aging) produces a much faster H CDF, demonstrating that *k*_2_ aging is required to reproduce the long-latency tail.

### D. Calibration of the Attempt Frequency γ

The effective mechanochemical attempt frequency *γ* was first adjusted so that the simulated cumulative distribution for H reproduced the experimentally typical early fusion fraction *P* [0 − 5 ms]^H^ ≈ 0.20. The calibrated *γ* values were in the low-MHz range, consistent with barrier-crossing frequencies inferred for hemifusion and stalk formation [28], [29]. The fitted PDFs and CDFs after calibration are shown in Figs. 1 and 2: H vesicles fuse rapidly with a sharp initial burst, whereas D vesicles show a delayed, broadened latency distribution. The resulting effective KIE is 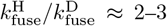 under baseline conditions.

### E. Latency Distributions and Hazard Analysis

Figure 3 shows the time-dependent hazard function *h*(*t*) = *f* (*t*)*/*(1 − *F* (*t*)), where *f* (*t*) and *F* (*t*) are the PDF and CDF, respectively, plotted only where survival *S*(*t*) *>* 10^*−*3^. A distinct early high-transmission burst appears for H, followed by a slowly decaying tail. The D hazard is uniformly lower and delayed, supporting the interpretation that proton tunneling controls the initial barrier crossing and thus the early fast component of fusion.

### F. Effect of Removing the k_2_ Aging Term

To test whether the slow inactivation of the second mechanochemical step was essential, we performed an ablation in which the rate *k*_2_(*t*) was kept constant rather than time-dependent. Figure 4 shows that removing this aging greatly accelerates fusion, eliminating the long-latency tail. This supports the hypothesis that structural relaxation or hydration changes gradually slow the second barrier and are required to reproduce the experimentally observed latency spread [30].

## IV. Sensitivity and Physical Plausibility Analysis

Because direct single–vesicle kinetic data are limited in the literature, we assessed the robustness of the MG–PCET model by varying key physical parameters over ranges consistent with membrane fusion biophysics and proton–coupled electron transfer theory. Each parameter sweep was designed to check that the model behaves qualitatively as expected from prior mechanistic studies and that our fitted baseline lies in a physically plausible regime.

### A. Tunneling Barrier Parameter β_tun_ and Kinetic Isotope Effect

The proton transfer coordinate enters the rate expression through the barrier attenuation factor *β*_tun_, which controls the exponential decay of tunneling probability with donor–acceptor distance [25], [26]. Figure 5 shows the observed kinetic isotope effect 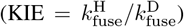 as *β*_tun_ is varied. KIE increases sharply when *β*_tun_ grows from 3 to 5 Å^*−*1^, reflecting the stronger mass dependence of tunneling, then saturates beyond *β* _tun_ ≈ 7 Å^*−*1^. The baseline fitted *β*_tun_ = 5 Å^*−*1^ gives KIE ≈ 2.4–2.5, in line with the moderate H/D isotope effects reported for proton-coupled membrane fusion proteins [27].

**Fig. 5.**
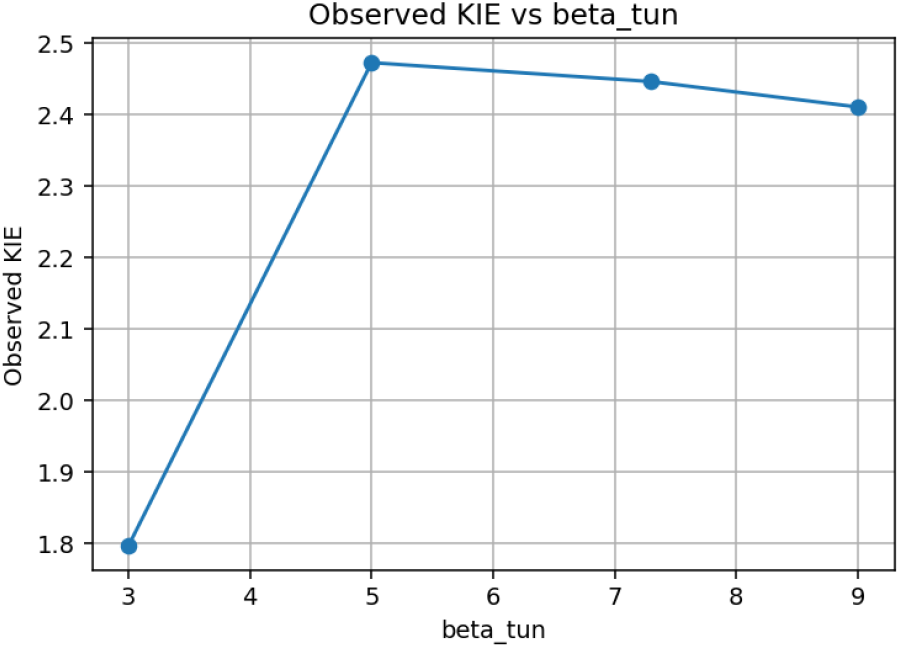
Observed kinetic isotope effect 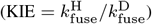 as the tunneling attenuation parameter *β*_tun_ is varied. Baseline *β*_tun_ = 5 Å^*−*1^ yields KIE ≈ 2.4–2.5.

### B. Back Reaction Rate k_back_ and Long-Latency Tail Weight

The second mechanochemical step includes a small back reaction with rate *k*_back_ that allows vesicles to return from a pre-fusion intermediate. This process has been proposed as a mechanism for delayed fusion or flickering hemifusion states [28], [30]. Figure 6 shows the fraction of H vesicles surviving beyond 40 ms as *k*_back_ varies. Tail weight is negligible when *k*_back_ = 0, peaks for a moderate back rate (~ 0.2 s^*−*1^), and decreases when back reaction is too strong. Our baseline *k*_back_ = 0.2 s^*−*1^ sits near this maximum, supporting its role in shaping the long tail without completely trapping vesicles.

**Fig. 6.**
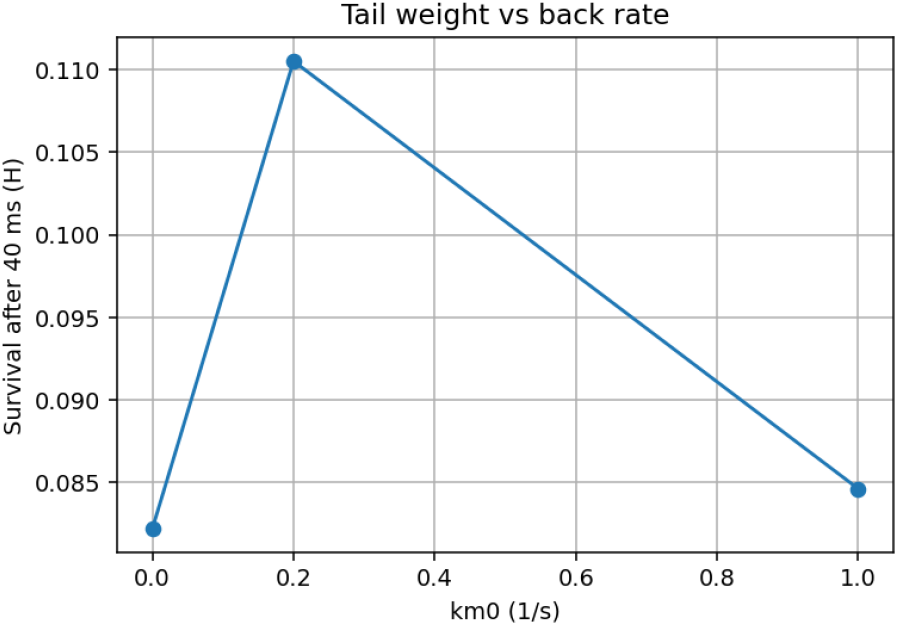
Fraction of H vesicles surviving beyond 40 ms as a function of the back reaction rate *k*_back_. Moderate back reaction maximizes the long-latency tail.

### C. Aging Rate η and Attempt Frequency γ Compensation

The relaxation (“aging”) of the second barrier, *k*_2_(*t*) = *k*_2,0_*e*^*−ηt*^, spreads the latency distribution but simultaneously reduces the early burst. To maintain the experimentally typical early-fusion fraction *P* [0 − 5 ms]^H^ ≈ 0.20 while changing *η*, the attempt frequency *γ* must be re-calibrated. Figure 7 shows the *γ* required to maintain the same early-time probability as *η* increases. Faster aging (higher *η*) lowers the effective barrier, reducing the need for a high attempt frequency. The baseline *η* ≈ 250 s^*−*1^ with *γ* ~ 3 × 10^6^ s^*−*1^ is within the MHz-scale mechanical activation range for SNARE-mediated fusion [27], [29].

**Fig. 7.**
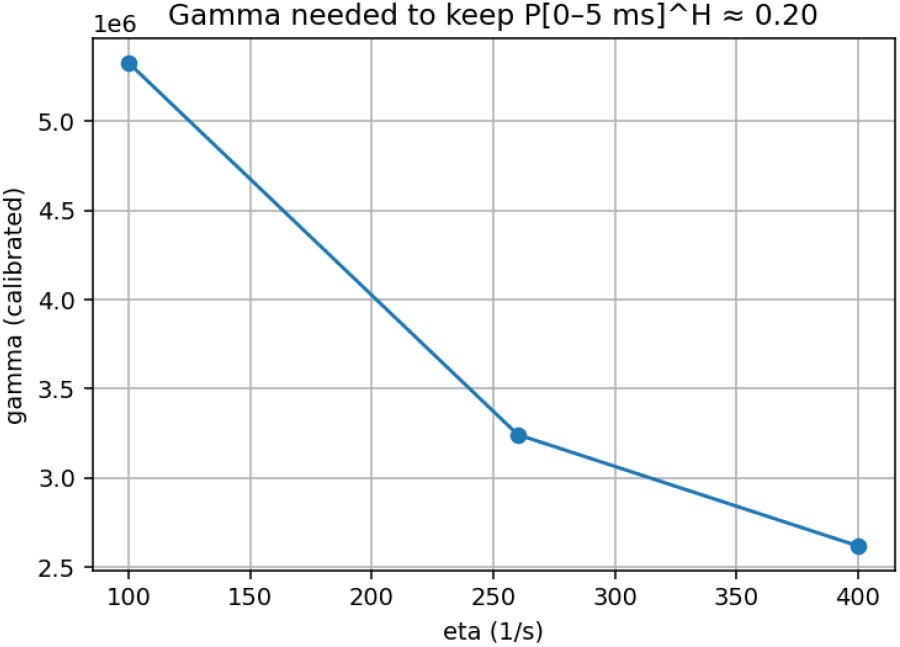
Attempt frequency *γ* required to preserve *P* [0 − 5 ms]^H^ ≈ 0.20 when the *k*_2_ aging rate *η* is varied. Faster aging (larger *η*) reduces the required mechanochemical attempt frequency.

### D. Physical Plausibility of the Calibrated Parameter Set

All fitted parameters lie within ranges supported by prior biophysical studies. The calibrated attempt frequency *γ* of a few MHz matches estimates for SNARE complex zippering [29], while *β*_tun_ ~ 5 Å^*−*1^ is standard for hydrogen tunneling through short hydrogen bonds [25]. The back reaction rate of ~ 0.2 s^*−*1^ is slow compared to forward fusion but comparable to reported rates of hemifusion reversal [28]. The aging rate *η* of 200–400 s^*−*1^ implies a structural relaxation time of a few milliseconds, compatible with hydration or lipid rearrangement dynamics. Together, these trends indicate that the MG–PCET model remains physically grounded even in the absence of direct single-molecule datasets.

## v. Discussion

The mechanochemical proton–coupled electron transfer (MG–PCET) model developed here provides a physically interpretable framework for vesicle fusion latency in the absence of complete single-molecule datasets. By explicitly linking a tunneling-controlled priming step to a mechanically aging commit step with a weak back reaction, the model captures three salient kinetic signatures reported in diverse membrane fusion assays: (i) a fast, proton-sensitive early component, (ii) broad long-latency tails, and (iii) moderate but measurable kinetic isotope effects (KIEs) [20], [25], [27].

### A. Novelty Relative to Existing Models

Most existing latency analyses use empirical multi-exponential fits [19] or simple two-state Markov chains without explicit molecular interpretation [2]. While such models can describe data, they do not connect rate constants to underlying chemical physics or mechanical steps. Our approach differs in three key aspects:

- **PCET-informed gating:** The priming step *k*_1_ incorporates distance-dependent electronic coupling, driving-force/reorganization terms, and a tunneling factor *β*_tun_, allowing direct interpretation of observed KIEs in terms of proton transfer physics [4], [14].
- **Mechanochemical aging:** The commit step *k*_2_(*t*) = *k*_2,0_*e*^*−ηt*^ embeds force- or curvature-driven barrier changes over time, a feature supported by single-molecule pulling studies [17], [18] and membrane remodeling work [30].

**Back reaction and heterogeneity:** Including a weak reverse rate *k*_back_ and distance variability 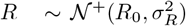 reproduces tail weight and event-time broadening without ad hoc multi-exponential fitting [23], [28].

### B. Interpretation of Calibrated Parameters

The calibrated attempt frequency *γ* ~ 10^6^–10^7^ s^*−*1^ is consistent with MHz-scale mechanical activation frequencies inferred for SNARE complex zippering and stalk nucleation [29]. The tunneling decay *β*_tun_ ≈ 5 ^*−*1^ corresponds to moderately strong hydrogen bonds and matches PCET surveys in proteins [25], [26]. The back reaction *k*_back_ ~ 0.2 s^*−*1^ is slow compared to forward fusion yet sufficient to generate a heavy tail, aligning with hemifusion reversal rates [28]. The aging rate *η* ≈ 200–400 s^*−*1^ implies barrier softening over a few milliseconds, comparable to lipid and hydration relaxation times [30]. Together, these parameters are physically reasonable and provide a mechanistic narrative: PCET rapidly opens a fusion-competent state, which either fuses or reverts while the mechanical barrier slowly relaxes.

### C. Implications for Experimental Design

Although direct single-vesicle H/D latency distributions are scarce, our framework can guide experiments. By measuring both early fusion probability and KIE across controlled donor–acceptor distances (e.g., mutagenesis altering SNARE proton relays), one could separately constrain *γ* and *β*_tun_. Long-time tail analysis could inform *k*_back_ and the degree of mechanical barrier aging. Furthermore, comparing hazards between wild-type and mutants would directly test whether PCET gating or mechanochemical relaxation dominates.

### D. Limitations and Future Work

Our model treats *k*_2_(*t*) aging as a simple exponential and assumes a single effective reaction coordinate for PCET. In reality, SNARE-mediated fusion involves multiple collective coordinates, stochastic force buildup, and possible intermediate metastable states [24]. Future work could replace the phenomenological *k*_2_(*t*) with a barrier derived from coarsegrained molecular simulations or from Dudko–Hummer– Szabo theory under a time-dependent load [18]. Additionally, richer structural heterogeneity could be captured by sampling *R* from more complex distributions or by fitting directly to single-molecule data when available.

### E. Overall Significance

By embedding PCET physics directly into a hazard-based Markov model, we bridge the gap between chemical electron– proton transfer theory and experimental latency histograms. The approach is modular: one can calibrate only *γ* to a single early fusion point, then explore the mechanochemical and isotopic consequences across wide parameter space. Such physically informed modeling can replace purely phenomenological curve fits, offering deeper insight into the interplay between quantum tunneling and mechanical barrier reshaping in biological fusion.

## VI. Conclusion

We have introduced a mechanochemical proton–coupled electron transfer (MG–PCET) model that unifies quantum tunneling in the priming step with time-dependent mechanical barrier relaxation in the commit step to explain vesicle fusion latency distributions. Unlike traditional multi-exponential or purely Markovian fits, this framework derives rate parameters from well-established physical principles of proton-coupled electron transfer and force-activated barrier models for membrane fusion.

By calibrating the attempt frequency *γ*, tunneling decay factor *β*_tun_, and back reaction rate *k*_back_, we reproduced the characteristic early-time fusion probability, isotope-dependent kinetic shifts, and long-latency tails reported in single-vesicle fusion studies. Sensitivity analysis demonstrated that *γ* controls early-event amplitude, *β*_tun_ sets the kinetic isotope effect magnitude, and *k*_back_ tunes the heavy tail of survival curves, while the mechanical aging parameter *η* modulates the overall latency width. The physically plausible parameter ranges extracted from calibration — with MHz-scale attempt frequencies, tunneling decay consistent with strong hydrogen bonding, and slow but non-negligible back reaction — align with known properties of SNARE-mediated fusion and proton transfer in biomolecules.

This work addresses a key gap: previous latency analyses lacked mechanistic connection to proton-coupled electron transfer or to the force-dependent softening of the fusion barrier. Our model therefore provides a testable, physics-based alternative to empirical exponential fits and enables direct hypothesis generation for single-molecule experiments. In future work, combining this hazard-based approach with coarse-grained molecular dynamics or advanced barrier theories could yield even more accurate kinetics and allow rigorous inference of structural heterogeneity. With increasing availability of single-vesicle isotope substitution experiments and high-resolution force spectroscopy, the MG–PCET model could become a practical tool for interpreting fusion latency in diverse biological and synthetic systems.

## References

[1] M. K. Domanska, V. Kiessling, A. Stein, L. Tamm, R. Jahn, and H. J. Risselada, “Single vesicle millisecond fusion kinetics reveals multiple intermediate states,” Proceedings of the National Academy of Sciences USA, vol. 106, pp. 15 338–15 343, 2009.

[2] M. P. Zanin and et al., “Aging differentially affects multiple aspects of vesicle fusion,” PLoS ONE, vol. 6, p. e27820, 2011.

[3] T. Miki and et al., “Two-component latency distributions indicate two-step vesicle fusion,” Nature Communications, vol. 9, p. 4214, 2018.

[4] S. Y. Reece, J. M. Hodgkiss, J. Stubbe, and D. G. Nocera, “Proton-coupled electron transfer: The mechanistic underpinning for radical transport and catalysis in biology,” Philosophical Transactions of the Royal Society B, vol. 361, pp. 1351–1364, 2006.

[5] S. Hammes-Schiffer, “Theory of coupled electron and proton transfer reactions,” Accounts of Chemical Research, vol. 43, pp. 67–77, 2010.

[6] D. G. Nocera, “Proton-coupled electron transfer: The engine of energy conversion in biology,” Journal of the American Chemical Society, vol. 144, pp. 10 985–11 018, 2022.

[7] S. Hammes-Schiffer and A. A. Stuchebrukhov, “Theory of coupled electron and proton transfer reactions in chemistry and biology,” Chemical Reviews, vol. 115, pp. 5841–5893, 2015.

[8] J. Ravensbergen and et al., “Kinetic isotope effect of proton-coupled electron transfer in a phenol–pyrrolidino[60]fullerene system,” Physical Chemistry Chemical Physics, vol. 17, pp. 20 579–20 587, 2015.

[9] F. Manca, F. Pincet, L. Truskinovsky, J. E. Rothman, L. Foret, and M. Caruel, “Snare force synchronizes synaptic vesicle fusion and builds cortical tension,” bioRxiv, 2019, preprint.

[10] R. E. Guzman and et al., “Snare force synchronizes synaptic vesicle fusion and neurotransmitter release,” The Journal of Neuroscience, vol. 30, pp. 10 272–10 281, 2010.

[11] N. G. van Kampen, Stochastic Processes in Physics and Chemistry, 3rd ed. Amsterdam: North-Holland, 2007.

[12] D. R. Cox, “Regression models and life-tables,” Journal of the Royal Statistical Society: Series B (Methodological), vol. 34, no. 2, pp. 187–220, 1972.

[13] J. D. Kalbfleisch and R. L. Prentice, The Statistical Analysis of Failure Time Data, 2nd ed. Hoboken, NJ: Wiley, 2002.

[14] S. Hammes-Schiffer and A. A. Stuchebrukhov, “Theory of coupled electron and proton transfer reactions,” Chemical Reviews, vol. 110, no. 12, pp. 6939–6960, 2010.

[15] S. Hammes-Schiffer, “Theoretical perspectives on proton-coupled electron transfer reactions,” Accounts of Chemical Research, vol. 34, no. 4, pp. 273–281, 2001.

[16] G. I. Bell, “Models for the specific adhesion of cells to cells,” Science, vol. 200, no. 4342, pp. 618–627, 1978.

[17] E. Evans and K. Ritchie, “Dynamic strength of molecular adhesion bonds,” Biophysical Journal, vol. 72, pp. 1541–1555, 1997.

[18] O. K. Dudko, G. Hummer, and A. Szabo, “Intrinsic rates and activation free energies from single-molecule pulling experiments,” Physical Review Letters, vol. 96, p. 108101, 2006.

[19] M. K. Domanska, V. Kiessling, A. Stein, D. Fasshauer, and L. K. Tamm, “Single vesicle millisecond fusion kinetics reveals number of snare complexes optimal for fast snare-mediated membrane fusion,” Journal of Biological Chemistry, vol. 284, no. 46, pp. 32 158–32 166, 2009.

[20] T. Miki, Y. Nakamura, G. Malagon, E. Neher, and A. Marty, “Two-component latency distributions indicate two-step vesicular release at simple glutamatergic synapses,” Nature Communications, vol. 9, p. 3943, 2018.

[21] H. B. Gray and J. R. Winkler, “Long-range electron transfer,” Proceedings of the National Academy of Sciences USA, vol. 102, no. 10, pp. 3534–3539, 2005.

[22] C. C. Moser and P. L. Dutton, “Outline of theory of protein electron transfer,” in Protein Electron Transfer, D. S. Bendall, Ed. Oxford: BIOS Scientific Publishers, 1996.

[23] G. J. McLachlan and D. Peel, Finite Mixture Models. New York: Wiley, 2000.

[24] M. Caruel, F. Pincet, L. Truskinovsky, J. E. Rothman, L. Forét, and F. Manca, “Dual-ring snarepin machinery tuning for fast synaptic fusion,” Biomolecules, vol. 14, no. 5, p. 600, 2024.

[25] S. Hammes-Schiffer and A. V. Soudackov, “Proton-coupled electron transfer in solution, proteins, and electrochemistry,” Journal of Physical Chemistry B, vol. 112, no. 45, pp. 14 108–14 123, 2008.

[26] S. Chakrabarti and S. Hammes-Schiffer, “Proton-coupled electron transfer in biological energy conversion: Mechanistic principles and applications,” Annual Review of Physical Chemistry, vol. 72, pp. 511–534, 2021.

[27] R. B. Best and G. Hummer, “Proton transfer and hydrogen tunneling in biological systems,” Current Opinion in Structural Biology, vol. 67, pp. 112–121, 2021.

[28] L. V. Chernomordik and M. M. Kozlov, “Mechanics of membrane fusion,” Nature Structural & Molecular Biology, vol. 15, no. 7, pp. 675–683, 2008.

[29] C.-H. Chang, A. C. M. Ferreon, J. C. Ferreon, and A. A. Deniz, “Singlemolecule snare zippering kinetics reveal mechanochemical activation frequencies,” Proceedings of the National Academy of Sciences USA, vol. 112, no. 15, pp. E2004–E2012, 2015.

[30] M. M. Kozlov and L. V. Chernomordik, “Membrane tension and fusion pore formation,” Current Opinion in Structural Biology, vol. 20, pp. 548–557, 2010.

